# SqueezeMeta, a highly portable, fully automatic metagenomic analysis pipeline

**DOI:** 10.1101/347559

**Authors:** Javier Tamames, Fernando Puente-Sánchez

## Abstract

The improvement of sequencing technologies has allowed the generalization of metagenomic sequencing, which has become a standard procedure for analysing the structure and functionality of microbiomes. The bioinformatic analysis of the sequencing results poses a challenge because it involves many different complex steps. SqueezeMeta is a full automatic pipeline for metagenomics/metatranscriptomics, covering all steps of the analysis. SqueezeMeta includes multi-metagenome support allowing the co-assembly of related metagenomes and the retrieval of individual genomes via binning procedures. SqueezeMeta features several unique characteristics: Co-assembly procedure or co-assembly of unlimited number of metagenomes via merging of individual assembled metagenomes, both with read mapping for estimation of the abundances of genes in each metagenome. It also includes binning and bin checking for retrieving individual genomes. Internal checks for the assembly and binning steps inform about the consistency of contigs and bins. Also, the results are stored in a mySQL database, where they can be easily exported and shared, and can be inspected anywhere using a flexible web interface allowing the easy creation of complex queries.

We illustrate the potential of SqueezeMeta by analyzing 32 gut metagenomes in a fully automatic way, allowing to retrieve several millions of genes and several hundreds of genomic bins.

One of the motivations in the development of SqueezeMeta was producing a software capable to run in small desktop computers, thus being amenable to all users and all settings. We were also able to co-assemble two of these metagenomes and complete the full analysis in less than one day using a simple laptop computer, illustrating the capacity of SqueezeMeta to run without high-performance computing infrastructure. SqueezeMeta is a complete system covering all steps in the analysis of metagenomes and metatranscriptomes, capable to work even in scarcity of computational resources. It is therefore adequate for in-situ, real time analysis of metagenomes produced by nanopore sequencing.

SqueezeMeta can be downloaded from https://github.com/jtamames/SqueezeMeta

## Introduction

The improvement of sequencing technologies has allowed the generalization of metagenomic sequencing, which has become a standard procedure for analysing the structure and functionality of microbiomes. Many novel bioinformatic tools and approaches have been developed to deal with the huge numbers of short read sequences produced by a metagenomic experiment. Aside from the simple overwhelming amount of data (currently in the range of terabases), a metagenomic analysis is a complex task comprising several non-standardized steps, involving different software tools which results are often not directly compatible.

Lately, the development of highly portable sequencers, especially those based on nanopore technologies (Deamer et al., 2016), has allowed the in-situ sequencing in scenarios where the necessity of obtaining quick results is paramount, for instance clinical scenarios of disease control or epidemics (Quick et al., 2015, 2016). Also, metagenomic sequencing has been done in-situ, for instance in oceanographic expeditions in the Antarctic ice (Johnson et al., 2017; Lim et al., 2014), illustrating the growing capability of producing sequences right away in the sampling campaigns. This will allow the informed planning of upcoming sampling experiments according to the results found in previous days. We foresee that this kind of application will be increasingly used in the near future. Therefore, the bioinformatic analysis should be done in a very short span of time (hours), and to be amenable to lightweight computing infrastructure.

A standard metagenomic pipeline involves read curation, assembly, gene prediction, and functional and taxonomic annotation of the resulting genes. Several pipelines have been created for automatizing most of these analyses (Abubucker et al., 2012; Arumugam et al., 2010; Eren et al., 2015; Glass and Meyer, 2011; Kim et al., 2016; Li, 2009). However, they differ in capacities and approaches. One of the most important differences is the necessity or not of the assembly step. Some of these platforms skip the assembly and consequently the gene prediction, and rely instead in the direct annotation of the raw reads. Nevertheless, there are several drawbacks of working with raw reads: Since it is based on homology searches for millions of sequences against huge reference databases, it usually requires very large CPU usage. Especially for taxonomic assignment, the reference database must be as complete as possible to minimize errors (Pignatelli et al., 2008). Besides, the sequences are often too short to produce accurate assignments (Carr and Borenstein, 2014; Wommack et al., 2008)

Assembly, on the contrary, is advisable because it can recover bigger fragments of genomes, often comprising many genes. Having the complete sequence of a gene and its context makes its functional and taxonomic assignment much easier and more trustable. The drawback of assembly is the formation of chimeras because of misassembling parts of different genomes, and the inability to assemble some of the reads, especially the ones from low-abundance species. The fraction of non-assembled reads depends of several factors, especially sequencing depth and microbiome diversity, but it is usually low (often below 20%). Recently, some tools have been developed to reassemble the portion of reads that were not assembled in first instance, increasing the performance of this step (Hitch and Creevey, 2018). Also, co-assembling related metagenomes can alleviate very much this problem, as we will illustrate in the results section.

Assembly is also advisable because it makes it possible the recovery of quasi-complete genomes via binning methods. The retrieval of genomes is a big step ahead in the study of a microbiome, since it allows to link organisms and functions, therefore contributing to a much accurate ecological description of the functioning of the community. It is possible, for instance, to determine the key members of the microbiome (involved in particularly important functions), to infer potential interactions between members (for instance, looking for metabolic complementation), and to advance in the understanding of the effect of ecological perturbations.

The best strategy for binning is the co-assembly of related metagenomes. By comparing the abundance and composition of the contigs in different samples, it is possible to determine which contigs belong to the same organism: these contigs having similar oligonucleotide composition, similar abundances in individual samples, and a co-varying pattern between different samples. In this way, it is possible to retrieve tens or hundreds of genomic bins with different levels of completion, that can be used as the starting point for a more in-depth analysis of the functioning of the microbiome.

SqueezeMeta is a fully automatic pipeline for metagenomics/metatranscriptomics, covering all steps of the analysis. It includes multi-metagenome support allowing the co-assembly of related metagenomes and the retrieval of individual genomes via binning procedures.

A comparison of the capabilities of SqueezeMeta and other pipelines is shown in Table 1. Most current pipelines do not include support for co-assembling and binning, while some allow importing external binning results to display the associated information.

**Table 1:**
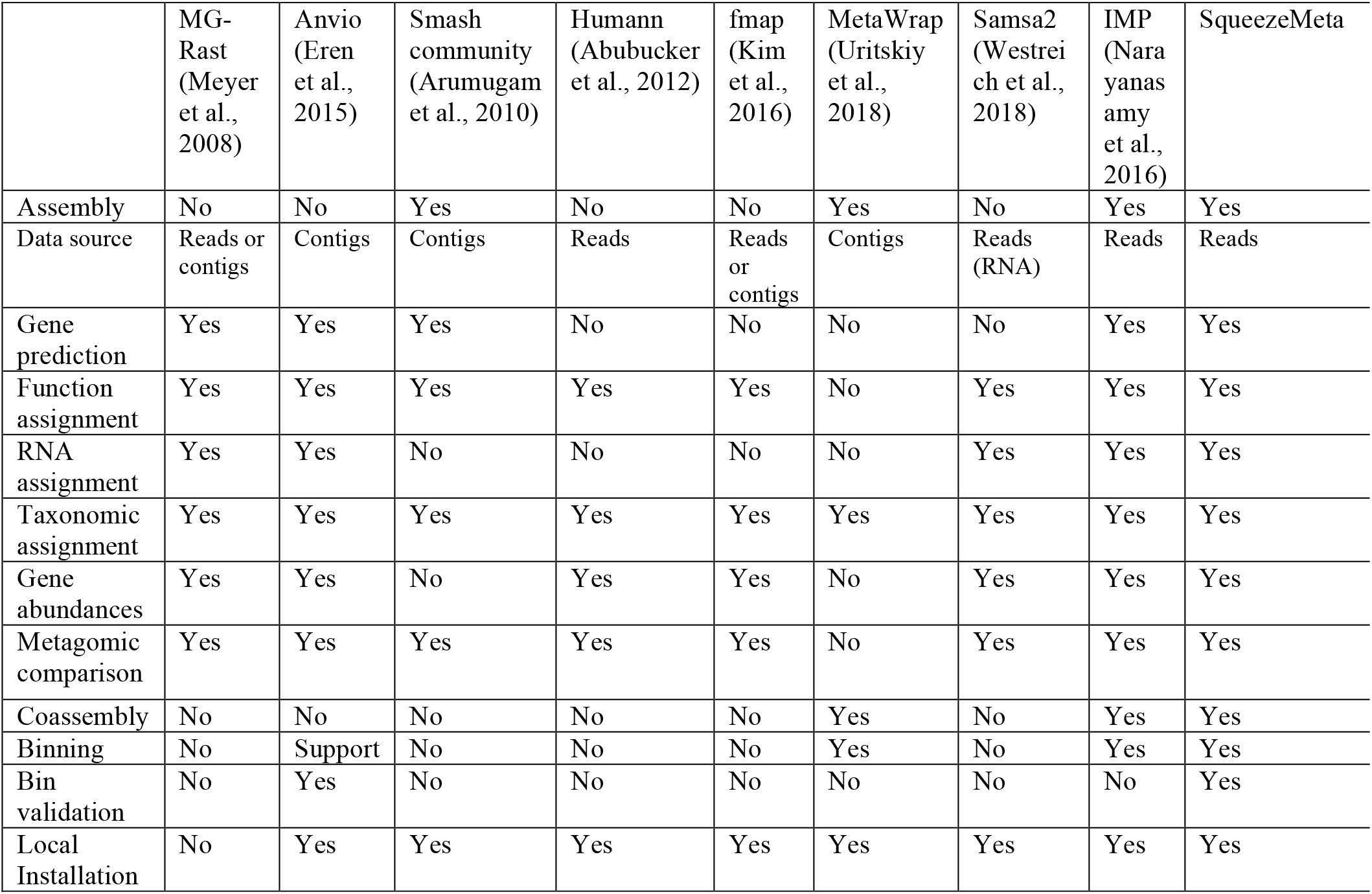
Features of different metagenomic analysis pipelines, in comparison to SqueezeMeta.

SqueezeMeta offers several advanced characteristics that make it different to existing ppipelines, for instance:

1. Co-assembly procedure coupled with read mapping for the estimation of the abundances of individual genes in each metagenome
2. An alternative co-assembly approach allowing to process an unlimited number of metagenomes via merging of individual metagenomes
3. Support for nanopore long reads
4. Binning and bin checking, for retrieving individual genomes
5. Internal checks for the taxonomic annotation of contigs and bins.
6. Metatranscriptomic support via mapping of cDNA reads against reference metagenomes, or via co-assembly of metagenomes and metatranscriptomes.
7. Included mySQL database for storing results, where they can be easily exported and shared, and can be inspected anywhere using a web interface

We have designed SqueezeMeta to be able to run in scarcity of computer resources, as expected for in-situ metagenomic sequencing experiments. By setting adequately all the components of the pipeline, we were able to analyse completely individual metagenomes, and even co-assemble related metagenomes using a desktop computer with only 16 Gb RAM. Also, the fully automatic nature of our system makes it very easy to use, not requiring any technical or bioinformatic knowledge.

SqueezeMeta can be downloaded from https://github.com/jtamames/SqueezeMeta

## Materials and Methods

SqueezeMeta is aimed to perform the analysis of several metagenomes in a single run. It can be run in three different modes (for a schematic workflow for the three modes, see Figure 1). These are:

-Sequential mode: All metagenomes are treated individually and analysed sequentially. This mode does not include binning, since each metagenome is treated independently.
-Coassembly mode: Reads from all samples are pooled, and a single assembly is performed. Then reads from individual samples are mapped back to the coassembly, allowing to obtain the coverage of contigs and of individual genes in these contigs. Based on these abundances, subsequent binning methods allow to classify contigs in genomic bins.
-Merged mode: The co-assembly is a very intensive process that requires plenty of computational resources, especially RAM memory. If the number of samples is high, the requirements can easily exceed the capabilities of the computing infrastructure. The merged mode of SqueezeMeta allows the co-assembly of a large number of samples, using a procedure similar to the one used in the analysis of TARA Oceans metagenomes (Tully et al., 2018). The samples are first assembled individually. The resulting sets of contigs are merged, by combining contigs with ≥99% semi-global identity, using CD-HIT (Fu et al., 2012). Then the remaining contigs are re-assembled using Minimus2 (Treangen et al., 2011) with parameters -D OVERLAP=100 MINID=95, to look for overlapping contigs coming from pieces of the same genome in different samples. The merging produces a single set of contigs, and the analysis proceeds as in the coassembly mode.

**Figure 1:**
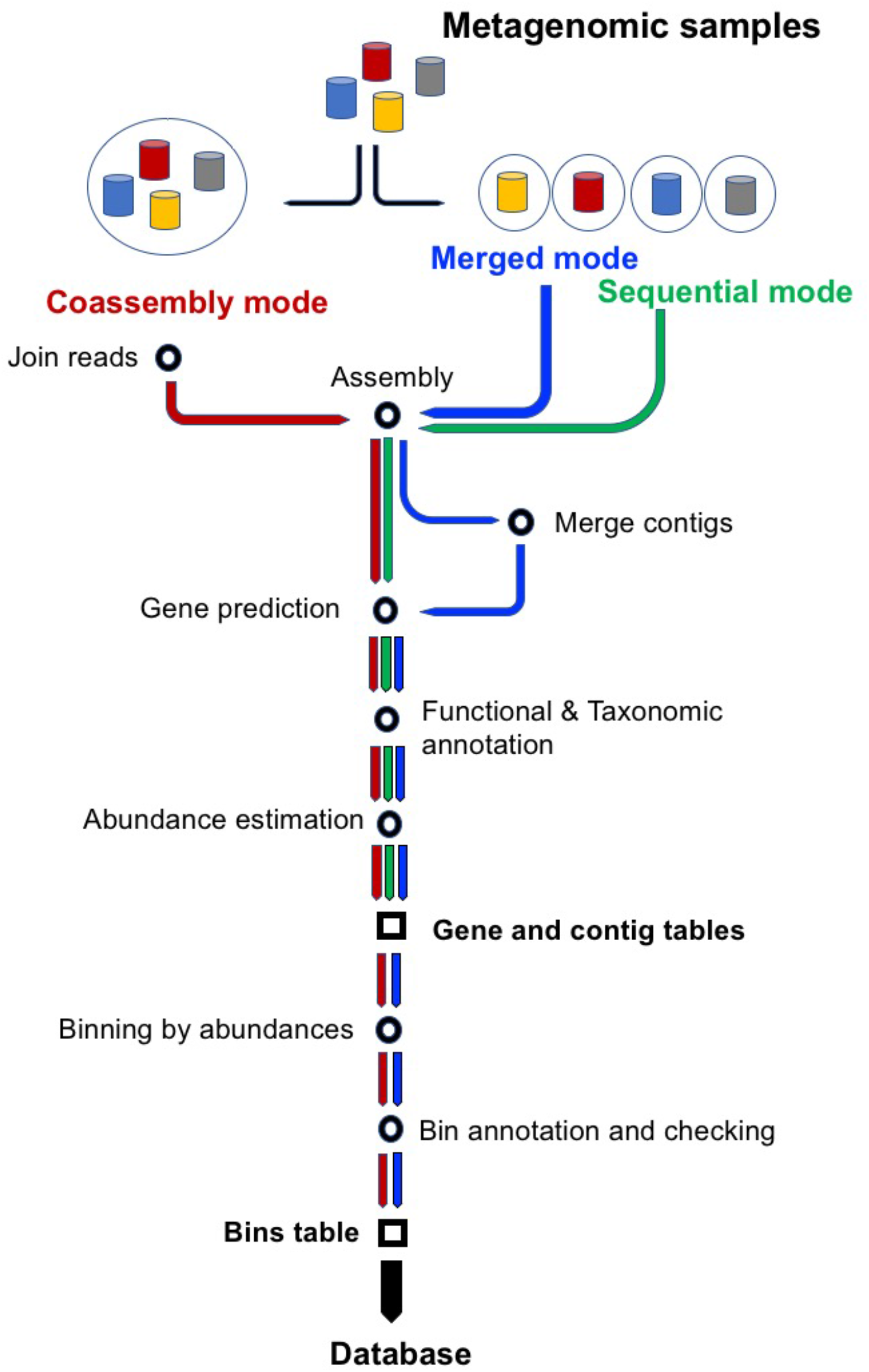
Workflow of the three modes of operation of SqueezeMeta: sequential, coassembly and merged. Starting from metagenomic samples, green, blue and red arrows indicate main steps in sequential, merged and coassembly modes. All modes create two of the three main tables of results: ORF and contig tables. Coassembly and merged modes also apply binning, and therefore they also create the bin table.

SqueezeMeta uses a combination of custom scripts and external software packages for the different steps of the analysis. A more detailed description of these steps follows:

Data preparation: A SqueezeMeta run is only requires a configuration file indicating the metagenomic samples and the location of their corresponding sequence files. The program creates the appropriate directories and prepares the data for further steps.
Trimming and filtering: SqueezeMeta uses Trimmomatic for adapter removal, trimming and filtering by quality, according to the parameters set by the user.
Assembly: When assembling big metagenomic datasets, computing resources, especially memory usage, are critical. SqueezeMeta uses Megahit (Li et al., 2015) as its reference assembler, since we find it has an optimal balance between performance and memory usage. SPAdes (Bankevich et al., 2012) is also supported. For assembly of the long, error-prone MinION reads, we use Canu (Koren et al., 2017). The user can select any of these assemblers. In the merged mode, each metagenome will be assembled separately and the resulting contigs will be merged and joined as described above. Either way, the resulting set of contigs is filtered by length using prinseq (Schmieder and Edwards, 2011), to discard short contigs if desired.
Gene and rRNA prediction: This step uses the prodigal software (Hyatt et al., 2010) to perform a gene prediction on the contigs, retrieving the corresponding amino acid sequences, and looks for rRNAs using barrnap (Seemann, 2014), classifying the resulting 16S rRNA sequences using the RDP classifier (Wang et al., 2007).
Homology searching: SqueezeMeta uses the Diamond software (Buchfink et al., 2015) for the comparison of the gene sequences against several taxonomic and functional databases, because of its optimal computation speed while maintaining sensitivity. Currently, three different Diamond runs are performed: against the GenBank nr database for taxonomic assignment, against the eggNOG database (Huerta-Cepas et al., 2016) for COG/NOG annotation, and against the latest publicly available version of KEGG database (Kanehisa and Goto, 2000) for KEGG ID annotation. It also classifies genes against Pfam database (Finn et al., 2014), using HMMER3 (Eddy, 2009). These databases are installed locally and updated at user’s request.
Taxonomic and functional assignment of genes: Custom scripts are used for this step of the analysis. For taxonomic assignment, SqueezeMeta implements a fast LCA algorithm that looks for the last common ancestor of the hits for each query gene using the results of the Diamond search against GenBank nr database (the most complete reference database available). For each query sequence, we select a range of hits having at least 80% of the bit-score of the best hit and differing in less than 10% of its identity percentage. The LCA is the taxon of lower rank that is common to most hits, since a small number of hits belonging to other taxa are allowed, to add resilience against, for instance, annotation errors. Importantly, our algorithm includes strict cut-off identity values for the diverse taxonomic ranks. This means that hits must pass a minimum aminoacid identity level in order to be used for assigning to a particular taxonomic rank. These thresholds are 85, 60, 55, 50, 46, 42 and 40% for species, genus, family, order, class, phylum and superkingdom ranks, respectively (Luo et al., 2014). Hits below these identity levels cannot be used to make assignments to the corresponding rank. For instance, a protein will not be assigned to species level if it has no hits above 85% identity. Also, a protein will remain unclassified if it has no hits above 40% identity. The inclusion of these thresholds guarantees that no assignments are done based on weak, inconclusive hits.
Functional assignments: Annotation of genes in COGs and KEGG IDs can be done using the classical best hit approach, or a more sensitive one considering the consistency of all hits (supplementary methods in Additional file 1). Briefly, the first hits exceeding a identity threshold for each COG or KEGG are selected. Their bitscores are averaged, and the ORF is assigned to the highest-scoring COG or KEGG whose score exceeds by 20% the score of any other, otherwise the gene remains unannotated. This procedure does not annotate conflicting genes with close similarities to more than one protein family.
Taxonomic assignment of contigs and disparity check: The taxonomic assignments of individual genes are used to produce consensus assignments for the contigs. A contig is annotated to the taxon to which most of their genes belong to (Additional file 1). The required percentage of genes assigned to that taxon can be set by the user, so that it is possible to accommodate missing or incorrect annotations of a few genes, recent HGT events, etc. A disparity score is computed for each contig, indicating how many of the genes are discordant with the consensus (Additional file 1). Contigs with high disparity could be flagged to be excluded from subsequent analyses.
Coverage and abundance estimation for genes and contigs: For estimating the abundance of each gene and each contig in each sample, SqueezeMeta relies on the mapping of the original reads onto the contigs resulting from the assembly. The software Bowtie2 (Langmead and Salzberg, 2012) is used for this task, but we also included Minimap2 (Li, 2018) for the mapping of long MinION reads. This is followed by Bedtools (Quinlan and Hall, 2010) for the extraction the raw number of reads and bases mapping to each gene and contig. Custom scripts are used to compute the average coverage and normalized RPKM values informing about gene and contig abundance. In sequential mode, SqueezeMeta would stop here. Any of the co-assembly modes allow to bin the contigs for delineating genomes.
Binning: Using the previously obtained contig coverage in different samples, SqueezeMeta uses different binning methods to separate contigs putatively coming from the same organism. Basically, binning algorithms classify contigs coming from the same genomes because their coverages co-vary along the samples, and their oligonucleotide composition is similar. Currently, Maxbin (Wu et al., 2015) and Metabat2 (Kang et al., 2015) are supported. Additionally, SqueezeMeta includes DAS Tool (Sieber et al., 2018) to merge the multiple binning results in just one set. SqueezeMeta calculates average coverage and RPKM values for the bins in the same way than above, mapping reads to the contigs belonging to the bin.
Taxonomic assignment of bins and consistency check: SqueezeMeta generates a consensus taxonomic assignment for the bins in the same way than it did for the contigs. A bin is annotated to the consensus taxon, that is, the taxon to which most of its contigs belong to. As previously, a disparity score is computed for each bin, indicating how many of the contigs are discordant with the consensus taxonomic assignment of the bin. This can be used as a first measure of the possible contamination of the bin.
Bin check: The goodness of the bins is estimated using the checkM software (Parks et al., 2015). Briefly, CheckM provides indications of the completeness, contamination and strain heterogeneity of a bin by creating a profile of single-copy, conserved genes for the given taxon and evaluating how many of these genes were found (completeness), and how many were single-copy (contamination and strain heterogeneity). SqueezeMeta automatizes CheckM runs for each bin, using the consensus annotation for the bin as the suggested taxonomic origin.
Merging of results: Finally, the system merges all these results and generates several tables: 1) a gene table, with all the information regarding genes (taxonomy, function, contig and bin origin, abundance in samples, aminoacid sequence). 2) A contig table, gathering all data for the contigs (taxonomy, bin affiliation, abundance in samples, disparity), and 3) A bin table with all information related to the bins (taxonomy, completeness, contamination, abundance in samples, disparity).
Database creation: These three tables and the optional metadata will be used in the creation of a MySQL database for the easy inspection of the data derived from the analysis. The database includes a web-based user interface that allows the easy creation of queries, so that the user does not need to have any knowledge on database usage to operate it (Figure 2). The interface allows queries on one table (genes, contigs or bins) or in combinations of tables, allowing complex questions such as “Retrieve contigs having genes related to trehalose from Bacteroidetes that are more abundant than 5x coverage in sample X” or “Retrieve antibiotic resistance genes active in one condition but not in other”. The resulting information can be exported to a table.

**Figure 2:**
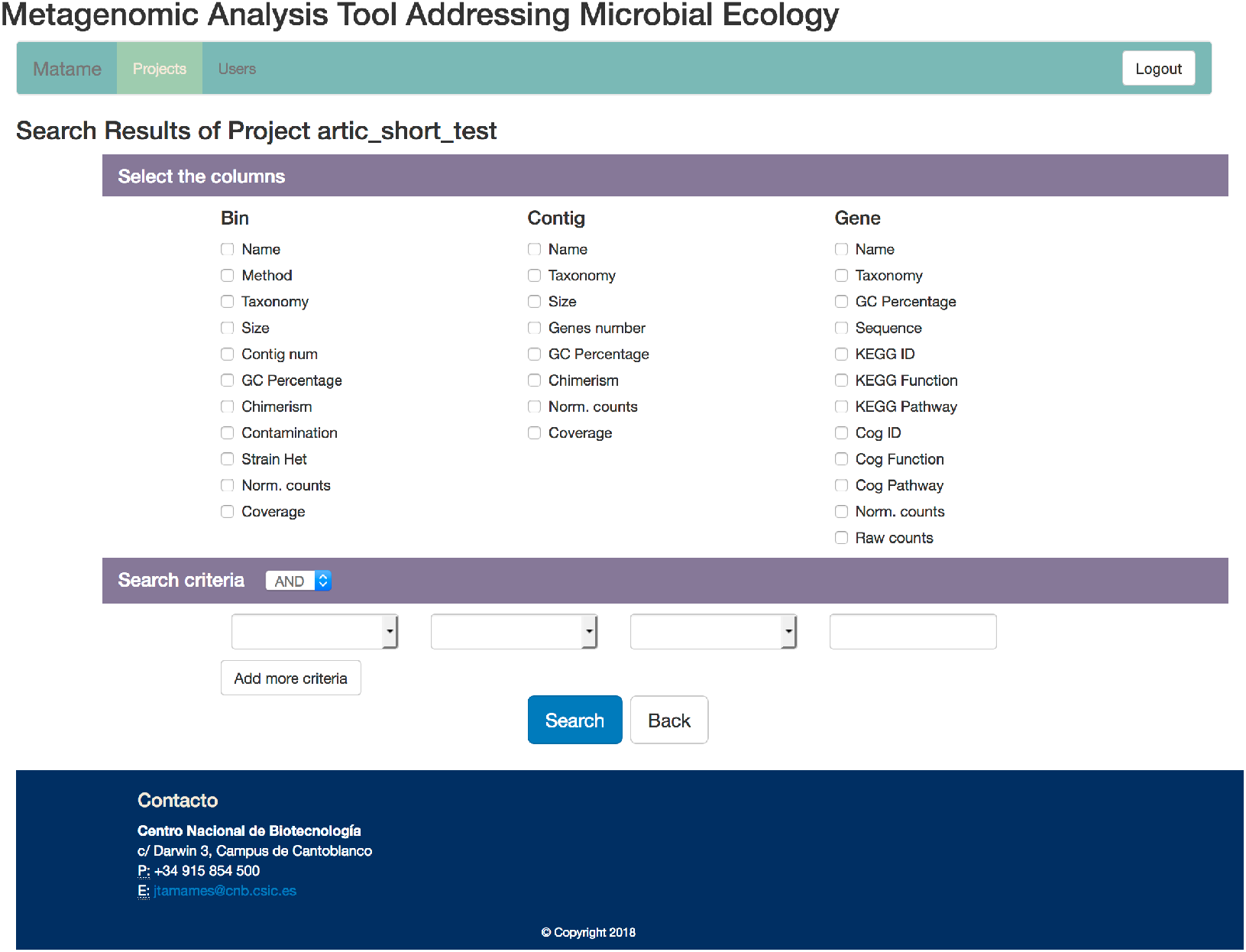
Snapshot of the SqueezeMeta user interface to its database A flexible and intuitive system for building queries allows to interrogate the database with complex questions involving combination of data from different tables.

When combining metagenomes and metatrancriptomes, the analysis of the latter can be done in a straightforward way by just mapping the cDNA reads against the reference metagenomes. In this way, we can obtain and compare the abundances of the same genes in both the metagenome and the metatranscriptome. But this will obviate these genes present only in the second, for instance genes belonging to rare species in the metagenome (therefore not assembled) that happen to be very active. SqueezeMeta can deal with this situation using the merged mode. Metagenomes and metatranscriptomes are assembled separately and then merged, so that contigs can come from DNA from the metagenome, cDNA from the metatranscriptome, or both. The normalization of read counts makes it possible to compare presence and expression values within or between different samples.

## Results

For illustrating the use of the SqueezeMeta software, we analysed 32 metagenomic samples corresponding to gut microbiomes of Hadza and Italian subjects (Rampelli et al., 2015), using the three modes of analysis. The total number of reads for all the metagenomes is 829.163.742. We used a 64-CPU computer cluster with 756 Gb RAM sited in the National Center for Biotechnology, Madrid, Spain. After discarding contigs below 200 bps, the total number of genes was 4.613.697, 2.401.848 and 2.230.717 for the sequential, merged and co-assembled modes, respectively. Notice that the number of genes is lower in the two latter modes that involve co-assembly since the genes that are present in more than one metagenome will be counted just once in the co-assembly (they are represented by just one contig product of the co-assembly) but more than one in the individual samples (they are present in one different contig per sample). A more accurate comparison is shown in Figure 3, where a gene in the co-assembly is assumed to be present in a given sample if it can recruit some reads from that sample. As coassemblies create a much bigger reference collection of contigs than individual metagenomes alone, even genes represented by a few reads in a sample can be identified by recruitment, while they will probably fail to assemble in the individual metagenome because of their low abundance. In other words, co-assembly will produce contigs and genes from abundant taxa in one or more samples, that can be used to identify the presence of the same genes in samples in which these original taxa are rare. Therefore, it allows to discover the presence of many more genes in each sample. The improvement of gene recovery for the smaller samples is also noticeable by the percentage of mapped reads. The individual assembly for small samples achieves barely 35% of read mapping to the assembled metagenome, indicating that most of the reads could not be used. The small size (and therefore low coverage) of the metagenome prevented these reads to be assembled. When co-assembling these samples with the rest, more that 85% of the reads could then be mapped to the reference metagenome, indicating that the co-assembly is able to capture most of the diversity found in these small samples.

**Figure 3:**
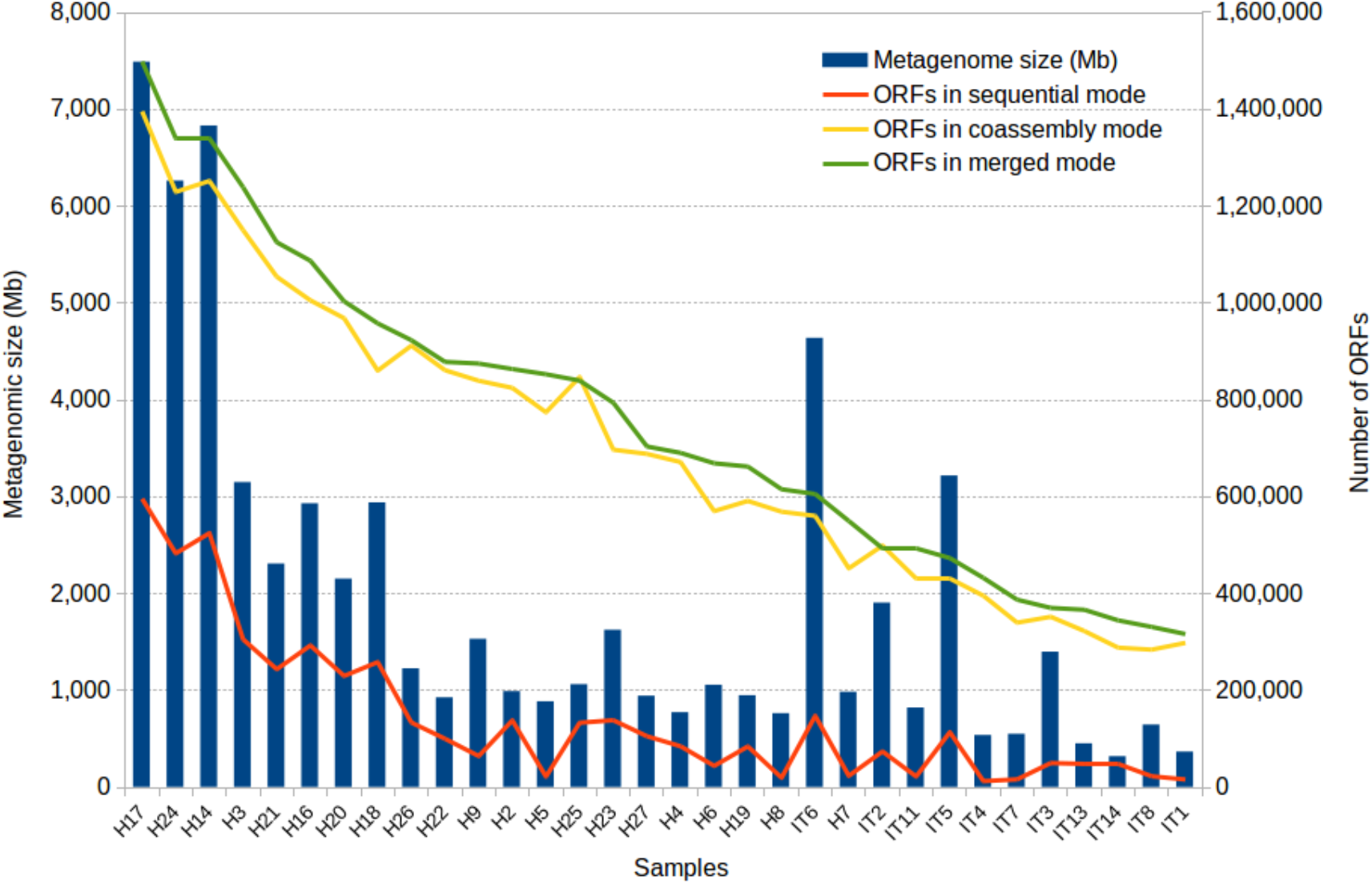
Results of the application of SqueezeMeta to 32 gut metagenomes of Hadza and Italian subjects. Size of Hadza and Italian metagenomes, and number of genes obtained using the three modes of analysis

Table 2 shows the characteristics of the analysis. Even if the merged mode obtains more contigs and genes than the co-assembly mode, we can see that the number of putatively inconsistent contigs (having genes annotated to different taxa) is lower in the second. Therefore, the co-assembly mode is more accurate than the merged mode, but the second has the advantage of being able to work with an almost unlimited number of metagenomes because of its lower requirements.

**Table 2:** Statistics on contigs and bins for the three modes on SqueezeMeta on Hadza & Italian metagenomes. Binning statistics refer to MaxBin results.

|  | Merged mode | Co-assembly mode | Sequential mode |
| --- | --- | --- | --- |
| Number of contigs | 893438 | 983350 | 2478560 |
| N50 | 3900 | 2357 | 2854 |
| Average perc of mapped reads | 85.01 | 89.47 | 74.47 |
| Contigs with phylum annotation | 719098 (80.4%) | 759903 (77.2%) | 1951445 (78.7%) |
| Contigs with disparity>0 | 6626 (0.7%) | 3772 (0.4%) | 7588 (0.3%) |
| Highly inconsistent | 4496 (0.5%) | 2993 (0.3%) | 5433 (0.2%) |
| contigs<br>(disparity>0.25) |  |  |  |
| Number of genes | 2401848 | 2230717 | 4613697 |
| Genes with COG<br>function | 1098635 (45.7%) | 982029 (44.0%) | 2164980 (46.9%) |
| Genes with KEGG<br>function | 835498 (34.8%) | 749892 (33.6%) | 1683636 (36.5%) |
| Total bins | 563 | 423 | N/A |
| Bins >90%<br>complete | 120 | 115 | N/A |
| Bins >50%<br>complete | 359 | 192 | N/A |
| High-quality bins<br>(>90% complete,<br><10% contam) | 50 | 67 | N/A |
| Good quality bins<br>>75% complete,<br><10% contam) | 82 | 112 | N/A |

The results of binning have been analysed according the values of completeness and contamination provided by CheckM (Table 3). Again, there are differences between the merged and the co-assembly modes, with the first providing more bins but less complete, and the second giving bins of higher quality. Both modes are capable to obtain quasi-complete genomes for tens of species, and hundreds of less complete genomes.

**Table 3:**
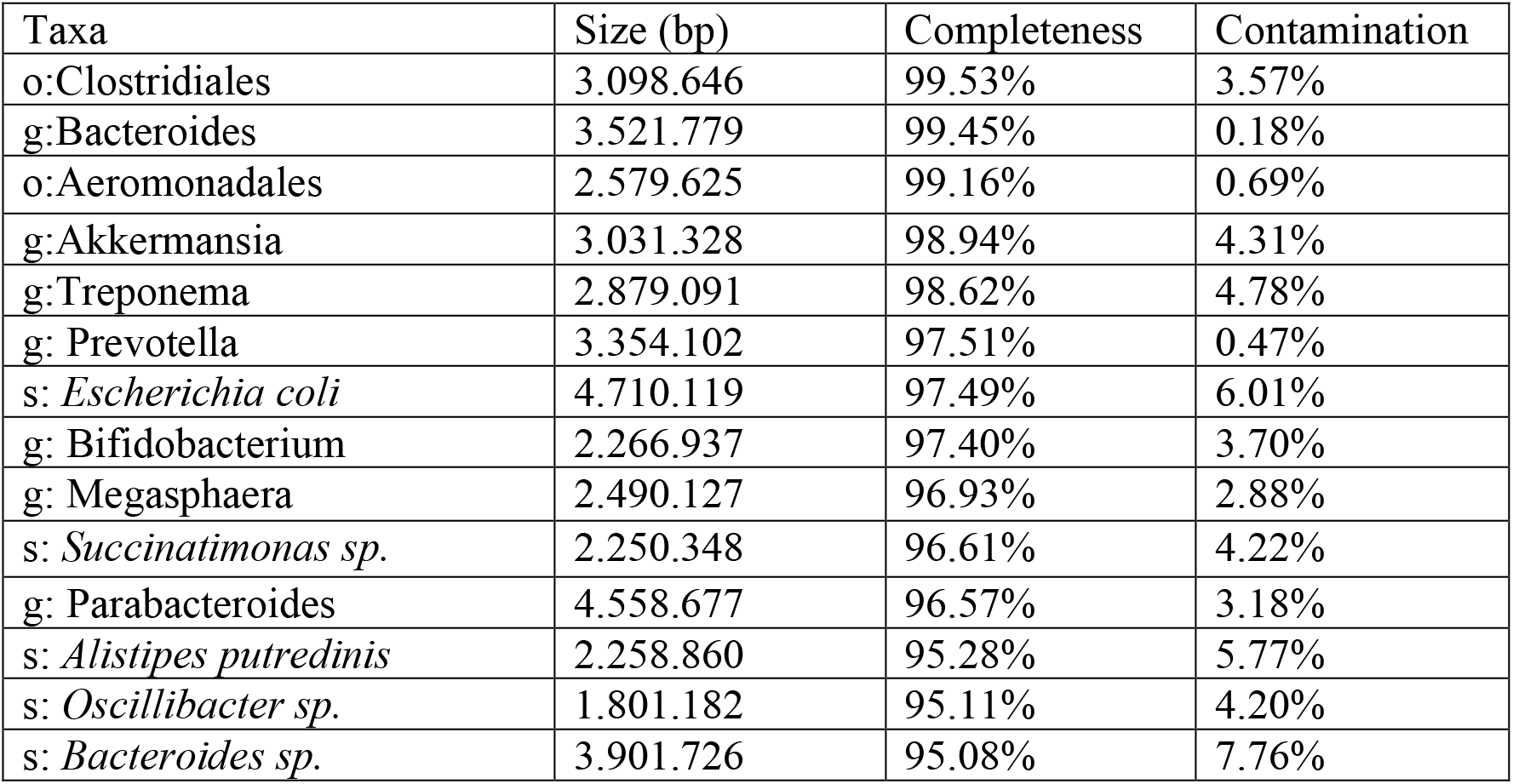
Example of some relevant high-quality bins (>90% completion, <10% contamination) obtained by the co-assembly mode of Hadza&Italian metagenomes. Taxa are labelled according to their taxonomic rank: o: Order; g: Genus; s: Species

Figure 4 shows the abundance distribution of bins in samples. The Italian subjects show a clear distinctive profile that make them clustering together. Bins belonging to the genera Bacteoides and Faecalibacterium are more abundant in these individuals than in Hadza individuals. The Hadza have increased diversity, and fall in different groups corresponding to the presence of diverse species, in accordance to the distinctions found using functional profiles (Rampelli et al., 2015). The microbiota of these individuals contains genera such as Allistipes or Prevotella that are not present in the Italian metagenomes. Also, Spirochaetes from the genera Treponema are only present in Hadza subjects, where are supposedly not associated to pathogenesis. This information is directly retrieved from SqueezeMeta results, and offers a revealing view of the genomic composition and differences between the samples. A similar result can be obtained for the functional annotations. The original functions represented in the bins can be used to infer the presence of metabolic pathways using the MinPath algorithm (Ye and Doak, 2009), that defines each pathway as an unstructured gene set and selects the fewest pathways that can explain the genes observed within each bin. The inference of several carbohydrate degradation pathways in the bins can be seen in Supplementary Figure 1.

**Figure 4:**
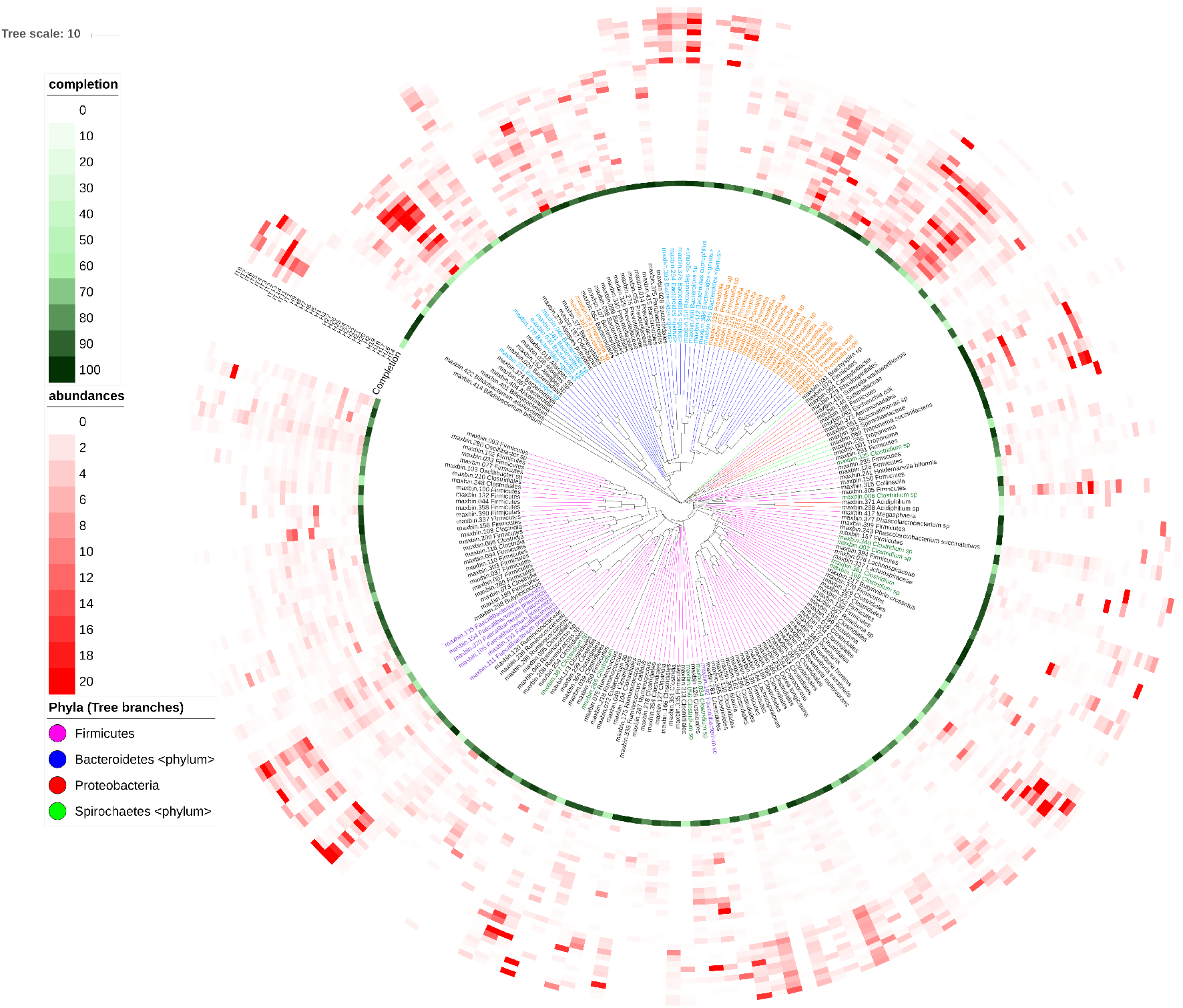
Abundance of bins in the diverse samples. Bins were compared with the compareM software (https://github.com/dparks1134/CompareM) to estimate its reciprocal similarities. The calculated distances between the bins were used to create a phylogenetic tree illustrating their relationships. The tree is shown in the inner part of the figure. The branches in the tree corresponding to the four more abundant phyla in the tree (Firmicutes, Bacteriodetes, Proteobacteria and Spirochaetes) were coloured. Bins were named with their id number and original genera, and the labels for the most abundant genera were also coloured. Outer circles correspond to: the completeness of the bins (green-coloured, most internal circle), and the abundance of each bin in each sample (red-coloured). Each circle correspond to a different sample (H, Hadza; I, Italians), and the red color intensities correspond to the abundance the bin in the sample. The picture was done using the iTOL software (https://itol.embl.de)

One of the motivations for the development of SqueezeMeta was making it capable to perform a complete metagenomic analysis on a limited computing infrastructure, like the one that can be expected in the course of in-situ metagenomic sequencing (Johnson et al., 2017; Lim et al., 2014). We created a setting mode (--lowmem) carefully tailored to run with limited amounts of resources, especially RAM memory. To test this capability, we were able to co-assemble two metagenomic samples from the Hadza metagenomes, composed of 40 millions of reads amounting almost 4Gb of DNA sequence. We ran the merged mode of SqueezeMeta using the --low-memory option in a standard desktop computer, using just 8 cores and 16 Gb RAM. The run completed in ten hours, generating 33.660 contigs in 38 bins and 124.065 functionally and taxonomically annotated genes. Using same settings, we also co-assembled ten MinION metagenomes from the gut microbiome sequencing of head and neck cancer patients (https://www.ncbi.nlm.nih.gov/bioproject/PRJNA493153), summing 581 Mb, in less than four hours. These experiments illustrate that SqueezeMeta can be run even with scarce computational resources, and it is adequate for its intended use of in-situ sequencing, where the metagenomes will be moderate in size.

## Discussion

SqueezeMeta is a highly versatile pipeline that allows to analyse a large number of metagenomes or metatranscriptomes in a very straightforward way. All steps of the analysis are included, starting with the assembly, subsequent taxonomic/functional assignment of the resulting genes, abundance estimation and binning to obtain as much as possible of the genomes in the samples. SqueezeMeta is designed to run in moderately-sized computational infrastructures, relieving the burden of co-assembling tens of metagenomes by using sequential metagenomic assembly and ulterior merging of resulting contigs. The software includes specific software and adjustments to be able to process MinION sequences.

The program includes several checks on the results, as the detection of possible inconsistent contigs and bins, and the estimation of completion of the latter using the checkM software. Finally, results can be easily inspected and managed since SqueezeMeta includes a built-in MySQL database that can be queried via a web-based interface, allowing the creation of complex queries in a very simple way.

One of the most remarkable features of this software is its capability to operate in limited computing infrastructure. We were able to analyse several metagenomes in a few hours using a virtual machine with just 16 Gb RAM. Therefore, SqueezeMeta is apt to be used in scenarios in which computing resources are limited, such as remote locations in the course of metagenomic sampling campaigns. Obviously, complex, sizeable metagenomes cannot be analysed with these limited resources, but the intended use of in-situ sequencing will likely produce a moderate and manageable size of data.

SqueezeMeta will be further expanded by the creation of new tools allowing in-depth analyses of the functions and metabolic pathways represented in the samples.

## Acknowledgements

This research was funded by projects CTM2016-80095-C2-1-R and CTM2016-80095-C2-1-R, Ministerio de Economía y Competitividad, Spain. This manuscript was made available as a pre-print at BioRxiv (Tamames and Puente-Sanchez, 2018)

We acknowledge Natalia García-García for helping in testing the system.

## Author Contributions

JT conceived and designed the tool. JT and FP created the software and performed all necessary testing. JT wrote the manuscript. Both authors read and approved the manuscript.

## Supporting information

**Additional file 1:** Description of novel algorithms implemented in SqueezeMeta.

**Supplementary figure 1:**
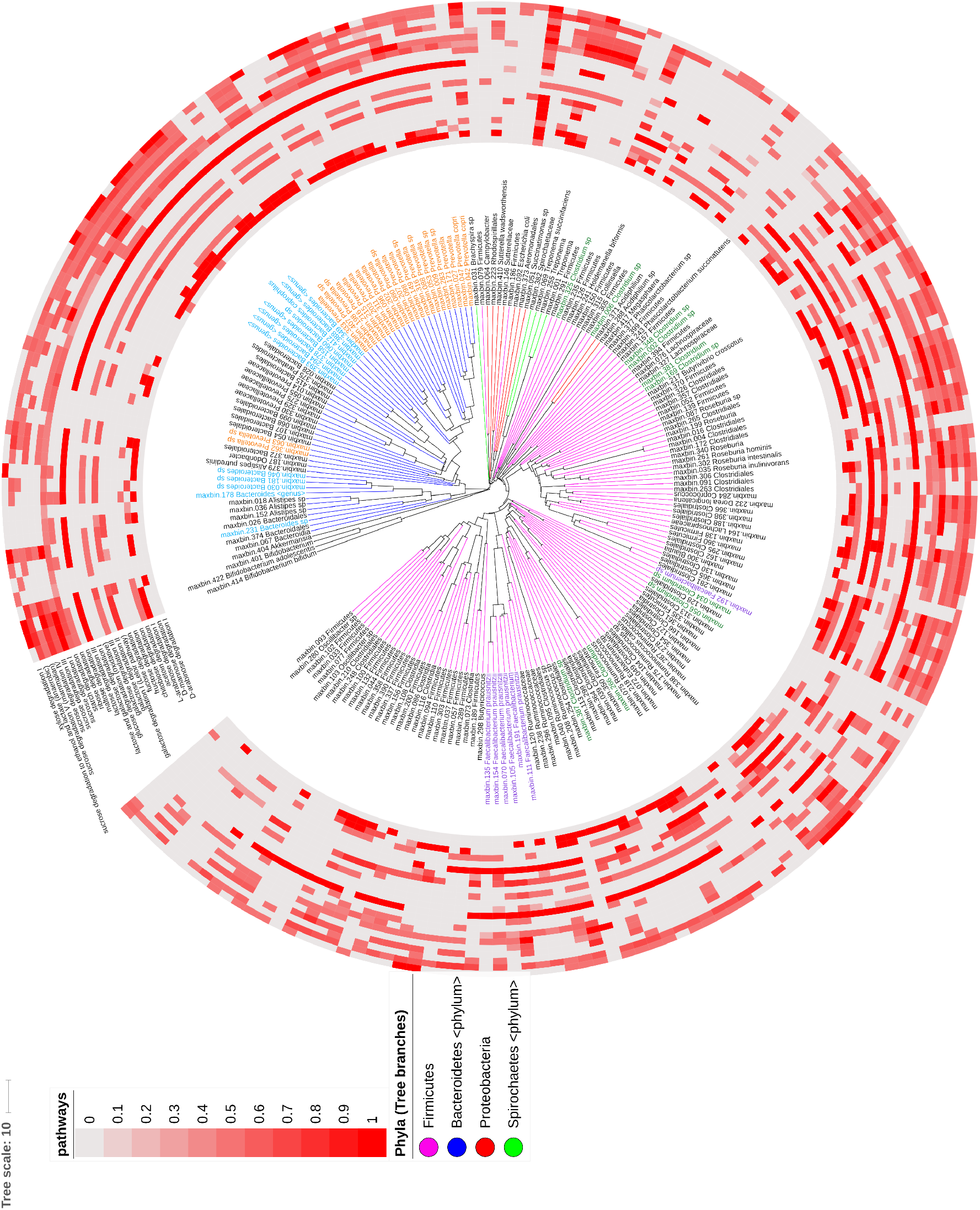
Presence of several carbohydrate degradation pathways in the bins. The outer circles indicate the percentage of genes from a pathway that are present in each of the bins. According to that gene profile, MinPath estimates if the pathway is present or not. Only pathways inferred to be present are coloured. As in figure 4, the tree of bins is done from a distance matrix of the amino acid identity of their orthoulogous genes, using the compareM software (https://github.com/dparks1134/CompareM). The four more abundant phyla are coloured (branches in the tree), as well as the most abundant genera (bin labels). The picture was done using the iTOL software (https://itol.embl.de)

